# Integrating Machine Learning with Musculoskeletal Simulation Improves OpenCap Video-Based Dynamics Estimation

**DOI:** 10.64898/2025.12.19.695562

**Authors:** Emily Y. Miller, Tian Tan, Antoine Falisse, Scott D. Uhlrich

## Abstract

**Objective:** Musculoskeletal dynamics influence the progression and rehabilitation of many movement-related conditions. However, accurately estimating whole-body dynamics using accessible tools, like smartphone video, remains challenging. Physics-based and machine learning (ML)–based dynamic predictions each offer advantages, but both approaches struggle to achieve both high accuracy and physical realism. Here, we created a hybrid ML–simulation framework to improve estimates of ground reaction forces, joint moments, and joint contact forces from smartphone video kinematics.

**Methods:** We used machine learning models to predict ground reaction forces and centers of pressure from video-based kinematics. The hybrid framework generates a dynamic simulation that tracks predicted forces and kinematics while enforcing dynamic consistency. We compared the hybrid model’s performance with a simulation-only approach and with ML forces applied through inverse dynamics. We evaluated mean absolute error from lab-based reference data (inverse dynamics from marker and force plate data) from 10 individuals walking.

**Results:** The hybrid model had 29% lower joint moment errors compared to simulations (p<0.001) and 45% lower errors compared to the ML-only approach (p<0.001). It also reduced vertical ground force error by 40% compared to simulations. The hybrid approach improved key metrics of joint loading related to knee osteoarthritis progression by 13–30% compared to simulations.

**Conclusion:** Our hybrid model outperforms purely physics-based and ML approaches for estimating dynamics from smartphone video during walking.

**Significance:** These methods move us closer to fast, accurate, and scalable assessments of whole-body musculoskeletal dynamics, which will enable large out-of-lab biomechanics studies and precision treatment of gait-related conditions.

## I. Introduction

Large-scale assessment of musculoskeletal dynamics could enable significant advances in injury risk prediction, personalized rehabilitation strategies, and outcomes evaluation. Measures of dynamics, such as ground reaction forces (GRFs), joint moments, muscle forces, and joint contact forces are essential for the study of neuro-musculoskeletal function. For example, joint moments are critical for many rehabilitation applications: they inform treatment planning in cerebral palsy [1], serve as surrogate measures of joint loading in osteoarthritis [2], [3], and provide control inputs for assistive exoskeletons [4], [5], [6]. Joint contact forces are similarly valuable for diverse applications: they directly quantify cartilage loading implicated in osteoarthritis progression [7], inform implant design in total joint replacement [8], and predict bone stress injury risk in athletes [9]. Despite their significance, routine measurement of 3D motion and dynamics remains uncommon in clinical practice. These measurements require complex, expensive lab-grade motion capture and force plate systems, which are costly, time-consuming, and impractical for regular clinical use [10], [11]. Consequently, high-fidelity biomechanical analyses are often restricted to lab-based research [12], [13], while clinical practice typically utilizes lower-fidelity functional metrics, such as the time required to walk 10 meters [14].

Recently developed approaches for estimating kinematics and dynamics with mobile sensors could lower the barrier to integrating biomechanical insights into clinical practice [10]. Of these emerging technologies, video-based motion capture has emerged as a promising alternative to marker-based motion capture. These techniques, powered by pose estimation models from computer vision, commonly use multiple videos to estimate 3D skeletal motion [15], [16]. Using this approach, we recently introduced the open-source OpenCap platform, which estimates lower-body kinematics using two smartphone cameras, achieving a mean absolute error of 4.5° across multiple activities compared to marker-based motion capture [17], [18]. OpenCap has since demonstrated robust performance outside the lab, enabling scalable kinematic analysis in clinical populations. In a study of 129 individuals with different subtypes of muscular dystrophy, OpenCap outperformed timed function tests in disease classification, capturing subtle kinematic differences, like variations in ankle height during the swing-phase of gait, that were missed by timing alone [19]. These applications underscore OpenCap’s potential for scalable, practical movement assessments outside of a motion capture laboratory.

From whole-body video-based kinematics, we can estimate dynamics using musculoskeletal modeling and physics-based simulation [20], [21], [22]. In laboratory environments, where both motion and force data are available, joint moments are typically calculated using inverse dynamics, which applies Newtonian mechanics to measured kinematics and external forces to determine joint moments. With video alone, however, ground reaction forces are not directly measured, making inverse dynamics infeasible during activities with both feet in contact with the ground. To address this, ground forces can be estimated in simulations by modeling foot-ground interaction using Hunt-Crossley contact spheres [22], where external forces on the foot are calculated based on the kinematics of the sphere relative to the ground plane. This results in a dynamically consistent (F-ma=0) set of kinematics and forces that track the input video-based kinematics. The simulation-based approach has been successfully demonstrated using kinematics derived from inertial measurement units [20] and from 3D video-based kinematics [17]. OpenCap has estimated kinetics using this approach with sufficient accuracy for several clinically relevant tasks, like estimating the knee adduction moment during walking and knee extension moment during chair rise [3], [17], [23], [24]. OpenCap can also detect intervention-induced changes in medial knee contact force, an important biomarker of osteoarthritis progression [25]. However, OpenCap’s performance varies across conditions, with reduced accuracy reported in several independent evaluations [26], [27]. Notably, while OpenCap detected changes in medial contact force, estimating its absolute magnitude remains challenging due to the complexity of resolving muscle forces from kinematics alone [23].

These physics-based simulations are valuable because they generate results that are both biomechanically plausible and dynamically consistent, meaning the predicted movements could be produced by the human musculoskeletal system and the external forces align with the observed accelerations. These simulations can also generalize across activities without requiring large training datasets. However, they are sensitive to assumptions in the model (e.g., body mass distribution, contact model parameters) and simulation process (e.g., objective function). For example, simulations that track lab-based motion capture data directly can modestly improve estimates of ground reaction forces, but substantial errors persist [18]. Thus, reduced accuracy in certain dynamic outcomes likely reflects limitations of the dynamic simulations, not solely errors in video-based kinematics.

Machine learning (ML) is a complementary approach for estimating musculoskeletal dynamics from kinematic data, bypassing the need for dynamic simulations. One ML strategy is to train models that directly predict joint moments from kinematic inputs [28], [29], [30]. While effective, this end-to-end approach must be retrained for each outcome and activity of interest, making it resource intensive. Alternatively, ML models can predict GRFs and centers of pressure (COPs) from video-based kinematics, which can then serve as inputs to traditional inverse dynamics analyses. This framework enables a similar workflow to estimating dynamic quantities, like joint moments and contact forces, as the lab-based approach. Foundation models like GaitDynamics [31], trained on large biomechanical datasets such as AddBiomechanics [32], exemplify the potential of this approach. On our OpenCap validation dataset (held out from model training), GaitDynamics predicted vertical GRFs with an average error of 3.9% body weight using marker-based kinematics, compared with 12.8% body weight error from physics-based simulations using video-derived kinematics [31]. Although GaitDynamics is neither trained nor tested with video inputs, it can likely improve the accuracy of ground reaction force estimates from video, compared to the OpenCap simulation approach. However, a key limitation of using ML-predicted forces is that they are likely dynamically inconsistent with the input kinematics (F-ma≠0), reducing the accuracy and biomechanical fidelity of downstream dynamic outputs, like joint moments or contact forces.

One approach to solving this is a hybrid model that leverages the accuracy of ML with the physical realism of biomechanical simulations. This would result in kinematics and dynamics that are both physically realistic and statistically likely, given large datasets of measured forces. Prior work has demonstrated that hybrid physics-based and ML modeling can improve kinematic accuracy with smaller training sets [7], generate training data from simulations [33], [34], embed physics into learning architectures [35], [36], and identify control strategies to reinforcement learning [37], [38]. Such a hybrid model could provide accurate, physically consistent GRFs and joint moments from smartphone video.

In this study, we present a hybrid framework that integrates ML predictions of GRFs and COPs into a physics-based simulation that tracks these values along with video-based kinematics. This approach aims to combine the accuracy of ML force-prediction models with the physical consistency of simulations to more accurately estimate musculoskeletal dynamics from video. We compared our hybrid approach to simulations tracking kinematics alone [17] and an approach that applies ML-predicted GRFs and COPs to video-based kinematics using inverse dynamics. We first compared GRF estimates between approaches, and hypothesized that the ML and hybrid approaches would more accurately estimate GRFs than the simulation-only approach. Additionally, we hypothesized that the hybrid approach would more accurately estimate (1) lower-extremity joint moments and (2) joint contact forces, compared to the simulation-only and ML-only approaches. Finally, to assess the clinical utility of our hybrid method, we evaluated the accuracy of clinically relevant knee loading measures associated with osteoarthritis progression, including medial contact force [25] and the knee adduction moment [3], [39].

## II. Methods

We evaluated the accuracy of four approaches for estimating ground reaction forces, joint moments, and joint contact forces from kinematics obtained from two smartphone videos processed with OpenCap, compared to lab-based motion capture and force plate analysis (Figure 1). Those approaches were: *Simulation*: dynamic muscle-driven simulation tracking kinematics [17], [22]; *ML*: machine learning prediction of GRFs and COPs via GaitDynamics [31] with inverse dynamics; *ML-Augmented*: the ML approach augmented with a new machine learning model that aims to more accurately predict COPs, with inverse dynamics; and *Hybrid*: dynamic simulations tracking kinematics, ground forces predicted by GaitDynamics, and COPs from our new model.

**Fig. 1.**
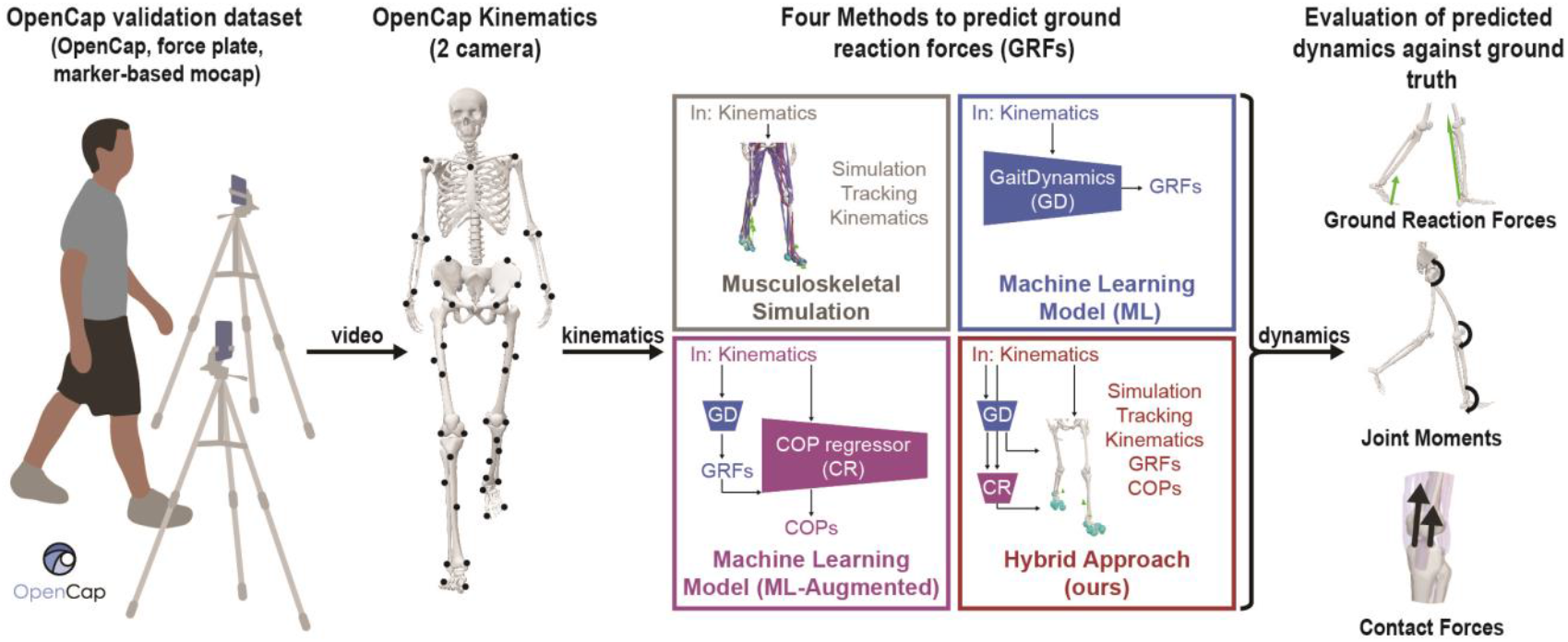
Overview of the machine learning and simulation-based methods to estimate musculoskeletal dynamics from video-based kinematics. Kinematics from two smartphone videos, processed with OpenCap, served as input to four approaches for estimating musculoskeletal dynamics: (1) simulation-only, which used muscle-driven optimal control simulations with Hunt–Crossley foot–ground contact; (2) machine learning, where GaitDynamics predicted ground reaction forces (GRFs) and centers of pressure (COPs) that were combined with video-based kinematics using inverse dynamics; (3) the same method as (2), except that a multilayer perceptron autoregressor was trained to predict the COP trajectories during stance; and (4) the hybrid framework, which tracked kinematics along with machine learning predictions of GRFs and COPs to balance physical consistency with improved external force estimation. All four approaches were compared against synchronously collected marker-based motion capture and force plate data. We evaluated accuracy on GRFs, joint moments, and joint contact forces.

### A. Experimental Data

We evaluated accuracy using the publicly available OpenCap dataset [17], which comprises video, marker-based motion capture, and force plate data from ten healthy adults walking over ground (age = 27.7±3.8, 6 female and 4 male, BMI = 22.9± 5.0 kg/m^2^) [14]. Video was captured using two smartphones positioned at approximately ±30° relative to the direction of walking, which we previously showed to provide equally accurate kinematics compared to three- and five-camera setups [17]. Each trial included synchronized marker-based motion capture (Motion Analysis Corp., Santa Rosa, CA, USA) and GRF data collected from three embedded force plates (Bertec Corp., Columbus, OH, USA). Each participant completed three walking trials with right, left, and right foot strikes on the three force plates. As input to our dynamics modeling approaches, we used the augmented anatomical marker positions derived by OpenCap [18], scaled OpenSim models [40], [41], and Inverse Kinematics results [42] from two OpenCap videos from this dataset.

### B. Simulation Approach

For the simulation-only approach, we used the simulation results from the original OpenCap publication; methods are described in detail there [17] and elsewhere [22]. Briefly, this approach generates muscle-driven tracking simulations of video-derived kinematics, formulated as an optimal control problem and solved using direct collocation. The simulations use a musculoskeletal model with 33 degrees of freedom [7], [40], [41], 80 lower-limb muscles and 13 ideal torque actuators for the lumbar, shoulder, and elbow degrees of freedom. Muscle-tendon dynamics are represented using Hill-type models with excitation–activation coupling [43], and skeletal motion is governed by Newtonian dynamics. Foot-ground interactions are modeled with six compliant Hunt-Crossley contact spheres per foot [18]. The optimal control problem minimized a composite cost function incorporating terms for actuator effort and tracking of generalized coordinates (i.e., joint angles and pelvis translations), velocities, and accelerations [44]. The optimal control problem is formulated using CasADi, derivatives are calculated using algorithmic differentiation [45], and it is solved using IPOPT [46]. Unlike machine learning methods, the simulation framework enforces dynamic consistency between kinematics and kinetics (i.e., no pelvis residual forces or moments). We extracted ground reaction forces and joint moments from the simulation results..

### C. Machine Learning Approach

For the ML approach, we used the GaitDynamics model to predict GRFs and COPs from the video-based kinematics [31]. GaitDynamics uses a transformer model, trained on the AddBiomechanics dataset [32], comprising 35 hours of synchronized motion and force data from 270 individuals across 15 studies. The model predicts GRFs and COPs from the kinematics of an OpenSim model [40].

Prior to predicting GRFs using GaitDynamics, we refined OpenCap kinematics to improve center-of-mass trajectories, which are essential for accurate GRF prediction. We optimized a vertical offset trajectory, applied to all marker data, to minimize vertical motion of heel and toe markers that were estimated to be in contact with the ground based on foot-marker velocity thresholds. Using the refined OpenCap kinematics, we estimated GRFs and COPs using the GaitDynamics model and joint moments using inverse dynamics.

To improve our machine learning predictions of COP, we trained a neural network to predict COP from kinematic and GRF data (i.e., “COP regressor” model; Figure 1). The model is a multi-layer perceptron with two hidden layers of 128 units each. We trained it using a marker-based motion capture and force plate dataset of 88 individuals walking overground at a self-selected speed (510 steps) [47]. Inputs are percentage of stance phase; vertical GRF (output from GaitDynamics); foot length; heel, toe, and midfoot positions and velocities; and the angle between a sagittal projection of a vector from heel to toe and the lab anterior direction. The model is autoregressive, incorporating a 16-sample COP history, resulting in a 27-dimensional input. We divided the dataset into 80% for training and 20% validation for architecture selection and hyperparameter tuning. To evaluate the impact of the new COP regressor, the ML-Augmented approach uses OpenCap kinematics, GaitDynamics GRFs, and COP regressor COPs, as inputs to inverse dynamics.

### D. Hybrid Simulation-Machine Learning Approach

The hybrid method combines OpenCap kinematics with ML-predicted GRFs and COPs using a dynamic simulation. It uses the same optimal control framework as the simulation-only approach [22], [41], [43], but uses a torque-driven model instead of muscle-driven one to improve both speed and convergence. Torque actuators included linear first-order activation dynamics [17], [22]. The simulation tracked OpenCap kinematics, ML-predicted GRFs from GaitDynamics, and ML-predicted COPs from the COP regressor. The simulations for two gait cycles of one individual did not converge, so we excluded them from analysis for all approaches. Unlike the simulation-only approach, we minimized rather than eliminated pelvis residual forces and moments. We chose this to balance convergence with the ability to track dynamically inconsistent input kinematics, GRFs, and COPs. The GRFs used for analysis were the resultant GRFs from the contact spheres in the simulation. We report the magnitude of pelvis residual forces and moments of all approaches.

### E. Joint Contact Force Estimation

To consistently compare joint contact forces across methods, we estimated muscle forces using a custom static optimization implementation [48] from the kinematics, GRF, and COP data from each approach. We computed joint contact forces using the Joint Reaction Analysis tool [42] and extracted compressive contact forces in the ankle, knee, and hip. We computed medial knee contact force from compressive force and internal adduction moment using the moment balance approach [49] and skin marker–based contact point estimates [50].

### F. Evaluation and Statistical Analysis

We evaluated the performance of all four methods against standard lab-based analyses using marker-based motion capture, force plate data, and inverse dynamics. We computed mean absolute error (MAE) for time series data over the left foot stance phase, the only stance phase with full bilateral measured ground forces. We computed MAEs for ground reaction forces, joint moments across 10 lower-extremity degrees of freedom (ankle dorsiflexion, subtalar supination, knee flexion, knee adduction, hip adduction, hip flexion, hip rotation, lumbar bending, lumbar extension, and lumbar rotation) [32], and joint contact forces (hip, ankle, knee). Additionally, we computed the magnitude of pelvis residual forces and moments. We normalize forces to body weight (BW) and moments to BW and height (BW*ht). For each metric, we computed an average MAE value for each participant, and we report the mean and standard deviation of MAEs across participants.

For statistical comparison, we compared the MAEs of GRFs, joint moments, and joint contact forces between the hybrid, ML, and simulation approaches. After confirming the normality of residuals using visual assessment of Q-Q plots and the Shapiro-Wilk test, we conducted pairwise t-tests on MAEs to test for differences in MAE among methods. To account for multiple comparisons and control the false discovery rate, we applied the Benjamini-Hochberg (FDR) correction to each of the three hypotheses independently (GRFs, joint moments, joint contact forces). We report corrected p-values with α=0.05. We limited statistical comparisons to our proposed method (hybrid) and the previously published benchmark models—simulation [17] and ML [31]. We did not statistically test the performance of the ML-Augmented method, since it is an intermediate analytical approach developed to isolate the specific contribution of improved COP estimation to overall model performance, rather than a competing model intended for validation.

## III. Results

### A. Ground Reaction Force Accuracy

The hybrid approach improved GRF estimation accuracy compared to simulation alone. For vertical GRF, the hybrid and ML models achieved average MAEs of 7.3–7.8 %BW (Figure 2), representing a 40–44% reduction from the 12.8 %BW error observed with the simulation-only approach (p<0.001). The hybrid model reduced medio-lateral GRF MAE by 51% compared to simulation (p<0.001), but there were not a significant difference in anterior–posterior GRF MAE (p=0.17). Notably, GRF errors from the ML model did not improve relative to simulation when using the original OpenCap kinematics (vertical GRF MAE=12.2 %BW; Figure S1, Supplemental Appendix); accuracy increased only when using kinematics with refined foot-floor and center-of-mass kinematics (Methods Section C).

**Fig. 2.**
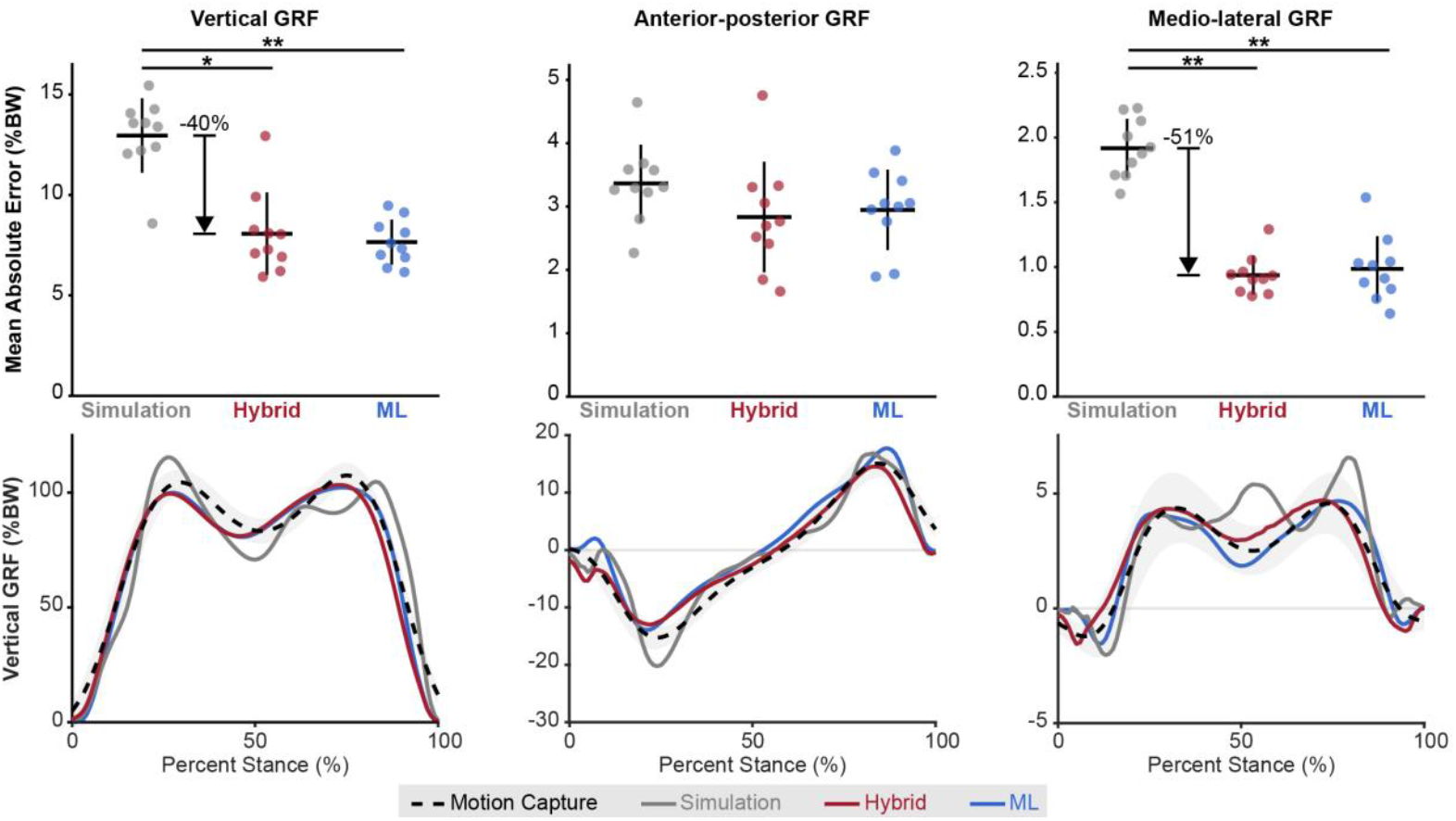
Ground Reaction Force Accuracy. Comparison of vertical, anterior-posterior, and medio-lateral ground reaction forces (GRFs) predicted by the three approaches, simulation-only, machine learning (ML), and the hybrid ML-simulation method (ours), against laboratory-based force plate measurements. Accuracy was evaluated using curve mean absolute error (MAE) during left foot stance. Lines indicate the mean across individuals. Error bars (top) and shading (bottom) represent standard deviations across participants. (*p<0.05; **p<0.001)

The COP regressor (ML-Augmented) achieved an average COP MAE of 1.8 cm on the training set and 1.4 cm on the held-out test set. When applied to the OpenCap dataset (held out from training), the average COP MAE was 1.5 cm.

### B. Joint Moment Accuracy

The hybrid approach had a joint moment MAE of 0.65 %BW*ht averaged across 10 lower-extremity degrees of freedom. This is 29% more accurate than the simulation-only approach and 45% more accurate than the ML approach (p<0.001; Figure 3). Additionally, the range of errors across degrees of freedom (Table 1) was lower for the hybrid approach (0.18-0.96 %BW*ht) compared to the simulation approach (0.21-1.40 %BW*ht). Although we did not test it statistically, the average MAE of the ML-Augmented model was between the ML and hybrid model (Figure 3). The degrees of freedom with the greatest moment MAE improvements from the simulation to the hybrid approach (Table 1) were ankle dorsiflexion (59%) and knee adduction (45%).

**TABLE I.** Joint Moment Accuracy During Walking.

| Joint | Hybrid MAE (%BW*height) | Simulations MAE (%BW*height) | Relative Improvement (%) |
| --- | --- | --- | --- |
| ankle dorsiflexion | 0.57 | 1.4 | 59 |
| hip adduction | 0.67 | 1.09 | 39 |
| hip flexion | 0.96 | 1.37 | 30 |
| hip rotation | 0.26 | 0.21 | -26 |
| knee flexion | 0.91 | 1.02 | 11 |
| knee adduction | 0.41 | 0.75 | 45 |
| lumbar bending | 0.39 | 0.63 | 39 |
| lumbar extension | 0.74 | 0.63 | -17 |
| lumbar rotation | 0.18 | 0.23 | 20 |
| subtalar supination | 0.36 | 0.26 | -37 |

\*Mean absolute error (MAE) for joint moment predictions during walking, reported in percent body weight times height (% BW\*height), comparing the hybrid method and simulation-only approach. Percent improvement shows the relative error reduction with the hybrid method compared to the simulation only approach. All moments were calculated during left stance. Positive values indicate improvement, while negative values reflect worse performance.

**Fig. 3.**
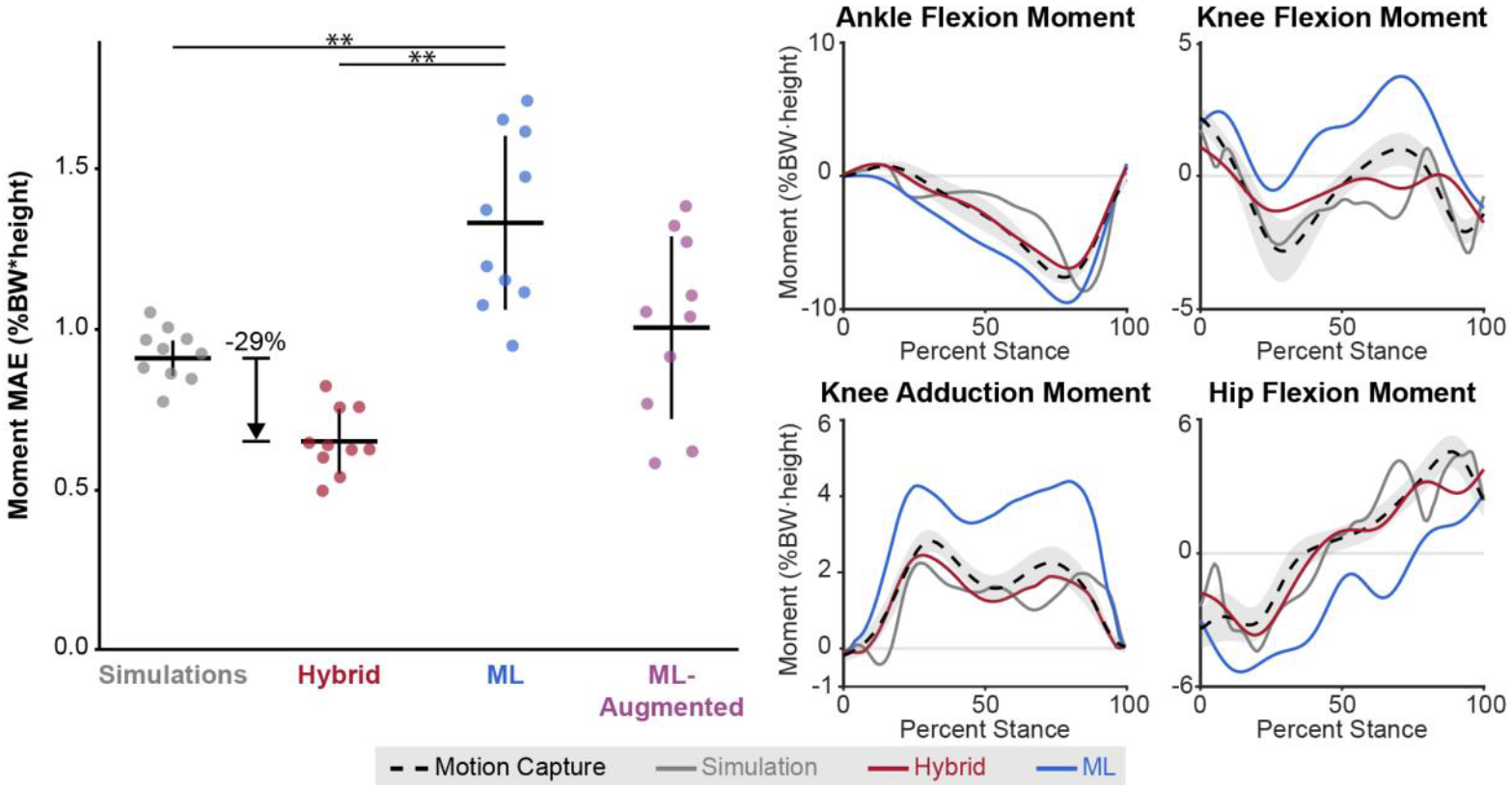
Joint Moment Accuracy. Comparison of joint moment predictions across 10 lower-limb degrees of freedom using simulation, machine learning (ML and ML-augmented), and the hybrid method (ours), with laboratory-based inverse dynamics as the reference standard. Accuracy was evaluated using curve mean absolute error (MAE) during stance and averaged across 10 lower-extremity degrees of freedom. Individual joint moment errors are shown in Table 1. Lines indicate the mean across individuals. Error bars (left) and shading (right) represent standard deviations across participants. (**p<0.001)

### C. Joint Contact Force Accuracy

The hybrid approach had the lowest MAE (0.92±0.09 BW) in joint contact force, averaged across the hip, knee, and ankle (Figure 4). This is a 13% improvement over the simulation approach (MAE=1.07±0.14 BW, p=0.039) and a 12% improvement over the ML approach (MAE=1.05±0.15 BW, p=0.040). Similar to joint moments, the ML-Augmented approach had an MAE between the hybrid and ML approach, though not tested statistically.

**Fig. 4.**
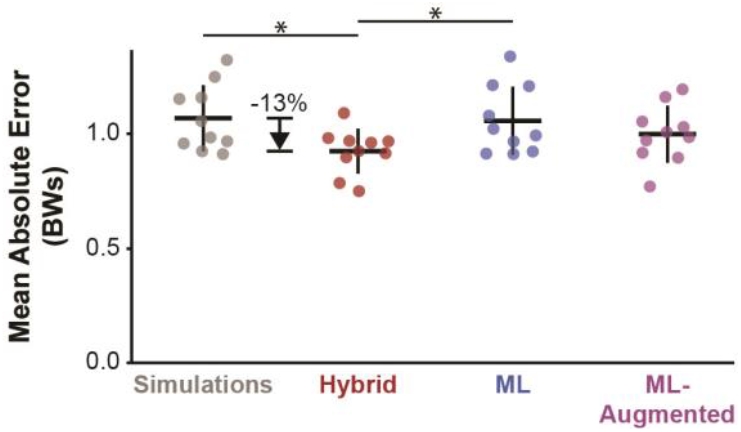
Joint Contact Force Accuracy. Average total contact force prediction error across four methods: simulation-only, hybrid ML–simulation, and two machine learning models, one using ground reaction forces (ML) and one using both GRFs and centers of pressure (ML-augmented). Error is shown as mean absolute error in body weights (BWs) during left stance. Error bars represent standard deviations across participants. Error bars represent standard deviations across participants. (*p<0.05)

Joint-level improvements were consistent across the lower limb (Table 2). Compared to simulations, the hybrid method reduced ankle contact force error by 13%, hip contact force error by 21%, and knee contact force error by 12%. The largest improvement was in medial knee contact force, where the hybrid approach reduced error by 26% compared to simulations.

**TABLE 2.** Accuracy of Joint Loading Measures during Walking.

| Joint | Hybrid MAE | Simulations MAE | Relative Improvement (%) |
| --- | --- | --- | --- |
| ankle contact force (BW) | 0.75 | 0.87 | 14 |
| hip contact force (BW) | 0.97 | 1.22 | 21 |
| knee contact force (BW) | 0.99 | 1.13 | 12 |
| Mean (BW) | 0.92 | 1.07 | 13 |
| **Knee loading metrics related to medial knee osteoarthritis progression** |  |  |  |
| medial knee contact force (curve) (BW) | 0.96 | 1.30 | 26 |
| first peak medial knee contact force (BW) | 1.01 | 1.98 | 49 |
| peak knee adduction moment (%BW*height) | 0.27 | 0.31 | 13 |
| knee adduction moment (impulse) (%BW*height) | 0.19 | 0.27 | 30 |

\*Mean absolute error (MAE) of joint contact forces normalized by body weight (BW), compared to marker-based motion capture and force plates. Positive improvement values indicate lower MAE in the hybrid vs. the simulation approach.

### D. Knee loading metrics

The hybrid approach improved accuracy of four key metrics associated with knee osteoarthritis, compared to simulations (Table 2). Accuracy improved by 30% (p=0.027) for the knee adduction moment impulse, by 49% (p=0.031) for the first peak of medial knee contact force, and by 26% (p=0.003) for the curve MAE of medial contact force. The MAE for the first peak knee adduction moment was not significantly different between methods (p=0.66); however, the hybrid approach’s 0.27 %BW*ht MAE is below clinically meaningful accuracy thresholds (0.5–2.2 %BW*ht) associated with the presence and progression of medial knee osteoarthritis [3], [54], [55], [56].

### E. Pelvis residuals

The average pelvis residual forces for the hybrid approach were 54.8N, which was 30% lower than the ML approach (Table 3). The simulation-only approach did not allow pelvis residuals (i.e., 0 residuals). Pelvis residual moments were 2.8 Nm for the hybrid approach, which was 93% lower than the ML approach. The hybrid approach produced larger residual forces than the motion capture and force plates (inverse dynamics), but smaller residual moments.

**TABLE 3.** Average pelvis residuals During stance.

|  | Hybrid (Nm) | Machine Learning (Nm) | Simulations (Nm) | Motion Capture (Nm) |
| --- | --- | --- | --- | --- |
| Residual Forces | $54.8 \pm 14.7$ | $77.8 \pm 18.8$ | $0.0 \pm 0.0$ | $28.2 \pm 6.0$ |
| Residual Moments | $2.8 \pm 1.2$ | $42.2 \pm 11.7$ | $0.0 \pm 0.0$ | $10.8 \pm 2.6$ |

\*Pelvis residuals were computed as the stance phase–averaged magnitude of the three-dimensional residual forces or moments (x, y, and z components) acting on the pelvis at each time point. Values are reported as mean ± standard deviation across subjects.

### F. Computation time

We ran all simulations and ML inference on a desktop with a 32-thread, 3.20 GHz central processing unit (Intel Core i9-14900KF) and a 24 GB graphics card (NVIDIA RTX 4090). Computing ground forces and joint moments for 1.3 seconds of walking data required 12.4 minutes for the simulation approach, 3.5 seconds for the ML approach, and 3.5 minutes for the hybrid approach.

## IV. Discussion

Our findings demonstrate the advantage of integrating machine learning with physics-based simulations to more accurately estimate musculoskeletal dynamics from video-based kinematics. Our hybrid ML-simulation framework outperformed both simulation-only and ML-only approaches, exhibiting the most accurate estimates of ground reaction forces, joint moments, and lower-extremity joint contact forces. Estimating the knee adduction below clinically meaningful accuracy thresholds and improving estimates of medial knee contact force and hip contact force illustrate the framework’s promise for advancing assessments of osteoarthritis progression and joint health. The ability to more accurately estimate dynamics from kinematics derived from mobile sensors addresses a key bottleneck in the field. Lab-based gold standards are accurate but cumbersome, and mobile sensing approaches often only measure kinematics. This work is a step towards estimating not only motion, but also clinically meaningful measures of musculoskeletal dynamics at scale using only smartphone videos.

The hybrid approach combines the strengths of ML and simulation, yielding estimates of musculoskeletal dynamics that are statistically likely and physically realistic. The simulation-only approach yields solutions that are dynamically consistent and biomechanically realistic. However, accuracy is limited by modeling and simulation assumptions that may not represent the individual, like model parameters or the neural objective function. Conversely, the GaitDynamics ML model predicted GRFs nearly twice as accurately as the simulations, but the GRFs and COPs were dynamically inconsistent with the input kinematics (i.e., large pelvis residuals). Thus, the ML-only joint moments were less accurate than the simulation-only approach, despite more accurate GRFs. The hybrid approach improved the dynamic consistency of the predicted GRFs and COPs with the kinematics, reducing pelvis residual forces and moments by 34–93% from the ML-only approach. The result was GRF accuracy similar to the ML-only approach, and joint moments that were 45% more accurate than simulation and ML-only approaches. The results align with prior research demonstrating the advantages of physics-informed machine learning methods for estimating dynamics from motion data [51], [52].

The improved center-of-mass kinematics and addition of the COP regressor model also contributed to the hybrid model’s accuracy. In OpenCap, global translations, estimated using the midpoint of the hip joints identified with computer vision algorithms, can be noisy. Refining these translations by minimizing foot motion during contact events improved pelvis kinematics and therefore GRF estimates: the GaitDynamics model did not improve vertical GRF estimates compared to simulations when using unrefined kinematics (Supplemental Figure S1) but improved them by 40% using refined kinematics (Figure 2). Additionally, joint moment accuracy was sensitive to both COP accuracy and dynamic consistency. Despite having substantially more accurate GRFs, the ML model, which did not enforce dynamic consistency, had less accurate joint moments than the simulation approach. Adding our COP regressor model improved both COP and joint moment accuracy over the ML model. Yet, the hybrid approach, by tracking GRFs from the GaitDynamics model and COPs from our COP regressor model in a more dynamically consistent and physiologically realistic way, had the most accurate joint moments, despite using identical input kinematics, GRFs, and COPs as the ML-Augmented approach, which combined them using inverse dynamics.

The hybrid method achieved clinically relevant accuracy levels across several kinetic outcomes. Vertical GRF errors are now smaller than minimal detectable change thresholds (MDCs of 10.2–13.0 %BW) [31], [53]. Additionally, improvements in vertical and medio-lateral GRFs likely improved frontal-plane kinetics, particularly the knee adduction moment and medial knee contact force. Notably, the hybrid model estimated the first peak knee adduction moment with MAE of 0.27 %BW*ht, well below the 0.5–2.2 %BW*ht clinical accuracy thresholds associated with medial knee osteoarthritis progression [3], [54], [55], [56]. This suggests that OpenCap, coupled with the hybrid dynamics approach, could help identify individuals with medial knee osteoarthritis who are at risk of rapid progression. This could support more targeted cohort selection in clinical trials and more informed treatment planning. In particular, clinically accessible tools for quantifying knee loading could be used to guide the personalized selection of biomechanical interventions for medial knee osteoarthritis, which can improve pain and slow cartilage degeneration when prescribed in a personalized manner [47], [57]. However, the accuracy improvement of the sagittal-plane knee moment (11%) was smaller than that of the frontal-plane knee moment (45%) and the sagittal-plane hip and ankle moments (30–59%). This may be because even small errors in ground contact, particularly in anterior–posterior force and COP location, or errors in the knee position can produce relatively large changes in the sagittal plane knee moment. Given the high prevalence of knee pathology and the link between the knee flexion/extension moment and knee contact force, improving its accuracy is an important direction for future work.

Although kinematic data can now be obtained from inertial sensors and video-based systems, kinetic measurements remain largely limited to laboratory environments with force plates. The growing availability of video recordings, from professional sports to clinical and residential settings, presents an opportunity to monitor biomechanics at scale, especially as single-camera methods continue to improve [58], [59]. Because kinetic variables more directly reflect tissue loading and injury mechanisms than kinematics alone, the ability to estimate dynamics from video could enable novel scientific questions— like estimating tissue loading during an injury from broadcast sports footage. It could also support large prospective studies of musculoskeletal function and enable more precise intervention selection and progress tracking in rehabilitation. Expanding kinetic measurement beyond the lab broadens our ability to understand loads in the musculoskeletal system over longer durations and in real-world contexts.

Several limitations should be noted. First, we validated on a dataset of young adults without mobility-related pathology. Although GaitDynamics has been tested in clinical populations, fine-tuning the model on pathology-specific datasets and applying our hybrid pipeline could likely improve accuracy for applications in specific clinical groups. Second, our work focused on gait, though the framework could extend to other activities using large, diverse movement datasets [28], [32]. Third, although our hybrid approach was four times faster than the simulation-only approach, it was still sixty times slower than the ML-only approach. For estimating ground reaction forces alone, the faster ML model is sufficient, but when joint moments are needed, the added biomechanical fidelity and associated computational time of the hybrid approach are likely justified.

Our goal is to make biomechanical assessment broadly accessible without the need for specialized equipment. To advance this vision, we integrated the hybrid modeling framework into OpenCap, an open-source platform that is freely available to the research community. Gait data can be collected in ~10 minutes using two smartphones, with kinematics automatically processed in the cloud [17]. We implemented the hybrid framework as companion software to the OpenCap data collection application. It segments gait cycles, runs ML inference, and generates physics-based simulations in an automated workflow. This combination of accessibility and biomechanical rigor enables high-fidelity estimates of movement and musculoskeletal forces in real-world settings, supporting wider integration of biomechanical insights into multi-disciplinary research and clinical practice.

## Conclusion

We introduced a hybrid machine learning–simulation framework that estimates important measures of musculoskeletal dynamics from smartphone video more accurately than existing machine learning or simulation methods. The approach combines the strengths of each method, producing statistically likely and biomechanically realistic estimates of dynamics, and it reached clinically relevant accuracy levels for several applications. By integrating our algorithms into the freely available OpenCap pipeline, this work could accelerate the use of biomechanical measures to improve the prevention and treatment of mobility-related disorders.

## Supporting information

Graphical Abstract

## Code and Data Availability

We used the publicly available OpenCap dataset available at https://simtk.org/projects/opencap. The source code is available at https://github.com/utahmobl/opencap-processing-grf and the software can be used through our web and mobile applications by visiting https://opencap.ai.

