## Supplementary figures and images for "Integrating Machine Learning with Musculoskeletal Simulation Improves OpenCap Video-Based Dynamics Estimation"

### Graphical Abstract

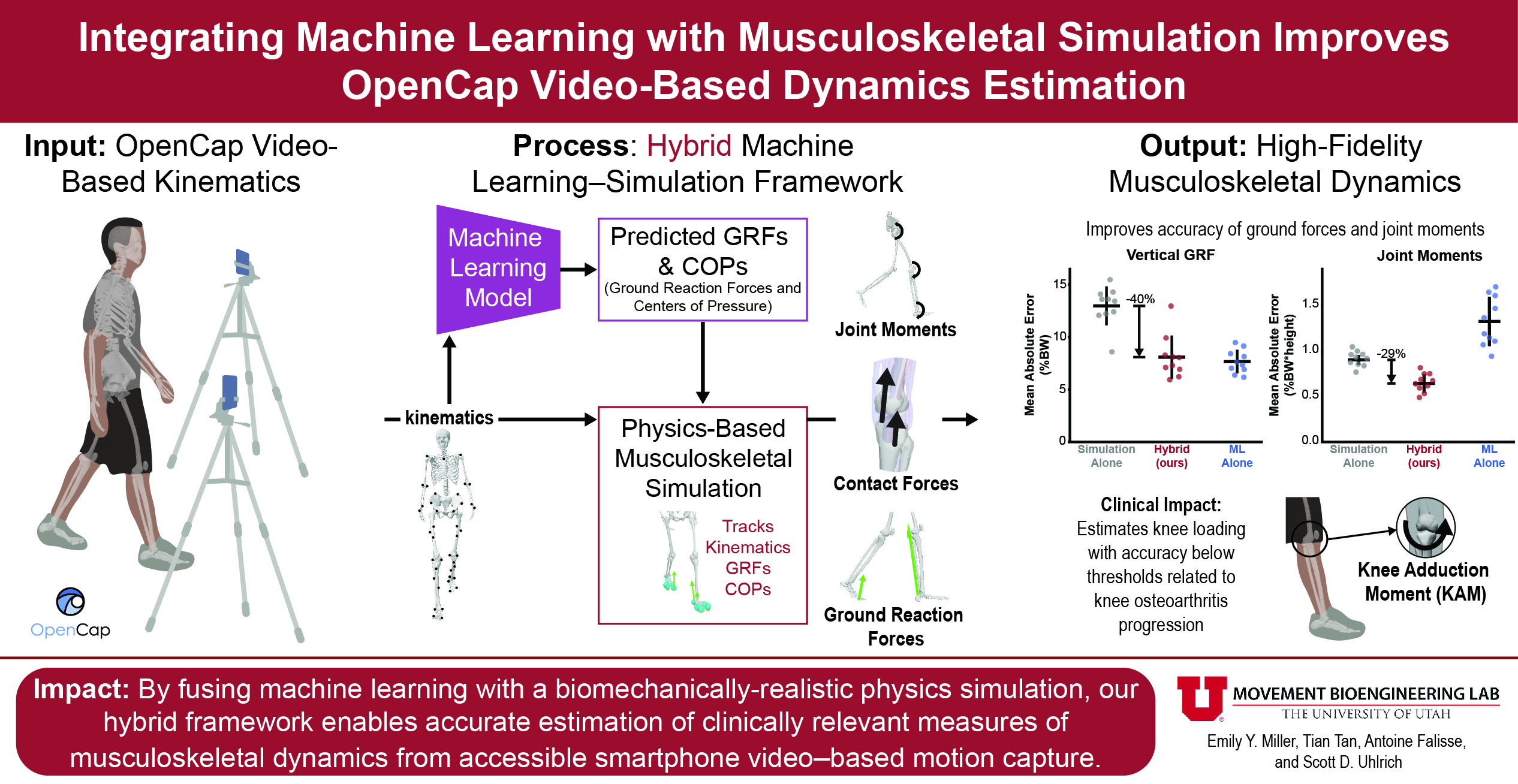
