## Supplemental Figure 1 for "Integrating Machine Learning with Musculoskeletal Simulation Improves OpenCap Video-Based Dynamics Estimation"

### Supplemental Material

**Refining center-of-mass kinematics improves ground reaction force prediction accuracy.** As shown in Figure S1, optimizing whole-body kinematics to minimize heel and toe motion during estimated contact periods substantially reduced vertical ground reaction force errors compared to physics-based simulation and the ML model applied to unrefined kinematics. This indicates that accurate ground reaction force estimation relies on realistic center-of-mass and foot kinematics.

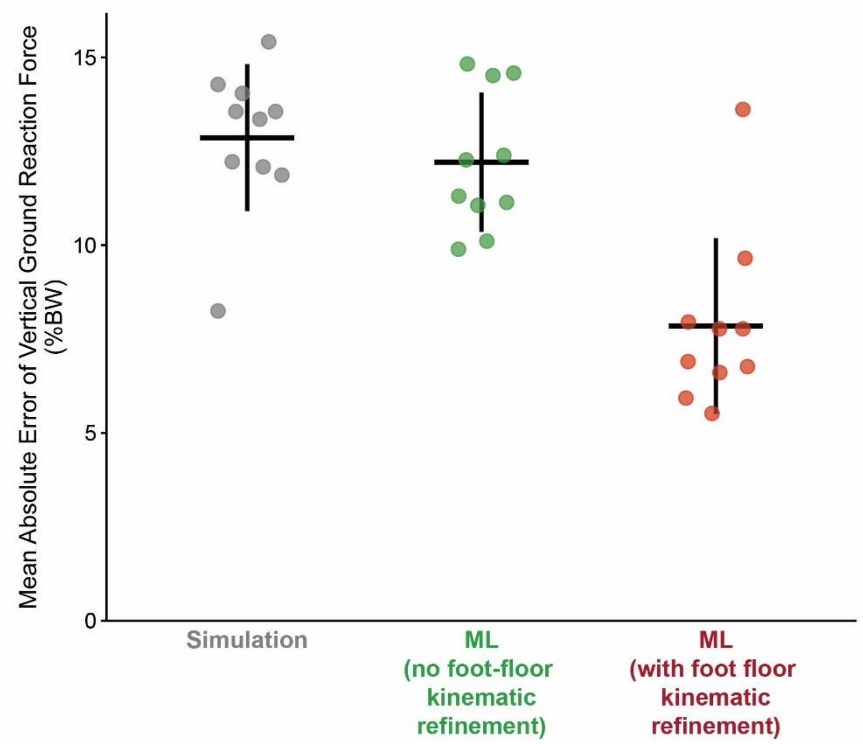


**Figure S1. Kinematic refinement center-of-mass trajectories improves vertical ground reaction force prediction accuracy.** Mean absolute error of vertical ground reaction forces is shown for three approaches, physics-based simulation, ML-only ground reaction force prediction (GaitDynamics model [1]), and the ML-only ground reaction force prediction with refined kinematics. Kinematic refinement minimized foot-floor motion during contact events, improving center-of-mass kinematics. The kinematic refinement procedure substantially improved vertical ground reaction force accuracy using the GaitDynamics model. Accuracy was computed across the full stance phase for all participants, and error bars indicate standard deviation across individuals (n=10).
